# Genome-wide SNP diversity in natural and cultivated populations informs restoration ecology with three *Calamagrostis* species from the Northwest Territories of Canada

**DOI:** 10.64898/2025.12.10.693508

**Authors:** Maria Jose Gomez Quijano, Yihan Wu, Claire Smith, Adriana Villalobos-Lopez, Pippa Secombe-Hett, Nathan Campbell, Zhengxin Sun, Robert I. Colautti

## Abstract

There is growing demand for data-driven frameworks to guide robust plant restoration strategies in response to anthropogenic disturbances. Several seed-sourcing (i.e., provenancing) strategies have been proposed, which balance the use of locally adapted genotypes against mixed genotypes to reduce mutation load or assistant migration to anticipate future climate scenarios. However, taxonomic uncertainty and lack of data characterizing genetic differentiation and gene flow have hindered provenancing strategies for many ecologically important non-model plant species—especially those in remote but vulnerable regions like the boreal forests of northern Canada. To guide provenancing strategies following anthropogenic disturbance in Canada’s Northwest Territories, we characterize species-specific markers, population structure, and hybridization among three *Calamagrostis* species. Double digest RAD sequencing (ddRAD) resulted in 2,951 polymorphic loci across 27 individuals, which we used to design loci for genotyping in thousands by sequencing (GT-seq), a cost-efficient target loci approach resulting in 256 polymorphic loci across 93 individuals from wild *C. canadensis*, *C. stricta* ssp. *inexpansa* and *C. purpurascens* populations. To help define the scale of ‘local’ populations for seed sourcing, we characterized geographic variation and population structure among 57 wild collection sites. We also assessed genetic relationships of wild *C*. *canadensis* to 69 individuals across eight commercially maintained populations used in restoration projects. We found that GT-seq yields similar genetic differentiation patterns as common neutral molecular marker approaches like ddRAD-seq. Specifically, we resolve morphologically misidentified individuals, identify genetic hybrids, and characterize the scale of genetic isolation-by-distance. Finally, we determined that three cultivar seed sources were genetically similar to southern wild populations, whereas five cultivars aligned with northern wild populations of *C. canadensis* in the Northwest Territories of Canada. Overall, our results highlight the benefits of cost-effective methods for genome-wide multi-locus genotyping to inform provenancing best-practices and support more effective and sustainable restoration efforts.

## 1. Introduction

Plants with broad geographical ranges often differentiate into genetically distinct populations with locally adapted phenotypes, thereby enhancing survival and reproduction throughout their range (Kawecki & Ebert, 2004; Leimu & Fischer, 2008; Hereford, 2009). The movement of genes among populations is an important factor affecting the degree of local adaptation. On one hand, when gene flow is limited and natural selection is strong, populations can undergo rapid genetic differentiation and local adaptation (Kawecki & Ebert, 2004; Aitken & Whitlock, 2013). On the other hand, populations with restricted gene flow are more prone to genetic drift, reducing genetic diversity and potentially limiting adaptive evolution and increasing the risk of local extinction (Lande, 1999; Frankham, 2005). Conversely, higher gene flow can introduce new alleles that accelerate the rate of adaptive evolution (Lowry & Willis, 2010; Aitken & Whitlock, 2013; Whiteley et al., 2015; Fitzpatrick & Reid, 2019). On the other hand, high gene flow can decrease population fitness by disrupting co-adapted gene complexes or swamping populations with maladaptive alleles, thereby disrupting the evolution of locally adapted genotypes (Slatkin, 1987; García-Ramos & Kirkpatrick, 1997; Todesco et al., 2016). The balance of gene flow with natural selection influences a broad range of potential outcomes, from the decline or recovery of rare endangered native species to the control or spread of invasive species (Kawecki & Ebert, 2004; Colautti et al., 2017; Wadgymar et al., 2022; Parmesan et al., 2023). As such, gene flow and population differentiation are important factors to consider in efforts to preserve, manage, and restore biodiversity in a rapidly changing world.

The relevance of local adaptation, genetic differentiation, and gene flow to management goals is particularly evident in the field of ecological restoration. A major goal for plant restoration is to improve rates of populations establishment and long-term persistence to accelerate the rate of habitat recovery and restore ecosystem function following anthropogenic disturbance (Lesica & Allendorf, 1999; Ruiz-Jaen & Mitchell Aide, 2005; Broadhurst et al., 2008; Kettenring et al., 2014). In particular, local provenancing—the selection of plant material based on their geographical origin—has been used to guide restoration practices based on an assumption that populations are locally adapted (Kiehl et al., 2010, 2014). However, rapid environmental change may result in maladapted populations if populations lag in their evolutionary response to novel optima. To address this problem, new approaches have been proposed that simulate gene flow to enhance standing genetic diversity by combining seeds from different source populations while matching for climate or other environmental variables. For example, composite (Broadhurst et al., 2008) and admixture (Breed et al., 2013) provenancing each combine seeds from local and non-local populations, albeit for different reasons—composite provenancing is designed to mimic natural rates of gene flow while admixture provenancing is designed to maximize heterosis. Regional admixture provenancing (Bucharova et al., 2019) sources materials from various local and non-local populations within an climatic and ecologically similar target region to preserve and maximize adaptation while increasing standing genetic variation relative to individual populations. Common to each of these strategies is the need to understand the spatial scale at which populations are “local” with respect to evolution. However, the degree of population genetic differentiation and gene flow in populations used for restoration purposes is often unknown, making it difficult to define the most appropriate spatial scale for provenancing and for predicting long-term persistence under future climatic conditions (Etterson & Shaw, 2001; Bucharova et al., 2019; Razgour et al., 2019; Maier et al., 2023).

Growing demand for seeds used for restoration projects has raised concerns that intensified cultivation and repeated harvesting could threaten the demographic stability of wild populations (Broadhurst et al., 2015; Pizza et al., 2021). On the other hand, sustainable harvesting and multi-generational propagation in cultivars are now common (Broadhurst et al., 2008; Kiehl et al., 2014; Espeland et al., 2017), but can increase rates of inbreeding, genetic drift, and unintentional selection, which compromise success of restoration projects (Bucharova et al., 2017; Conrady et al., 2023; Nevill et al., 2018; Pedrini et al., 2020; Pizza et al., 2021). Genetic screening can help identify the scale of ‘local’ populations and inform breeding programs to balance standing genetic diversity with local adaptation as the demand for cultivated seeds for restoration plans grow.

Demand for restoration projects is elevated in regions of high resource extraction, which can cause habitat fragmentation, biodiversity loss, soil degradation, water pollution, and community displacements (Haddad et al., 2015; Weiskopf et al., 2020; Arneth et al., 2021). For example, resource extraction (e.g., lumber, oil, minerals) in Canada’s Northwest Territories has contributed significantly to economic gain (NWT State of the Environment Report, 2022), but also contributes to climate change and impacts sensitive habitats through landscape changes, ocean acidification, permafrost degradation and changes in plant and animal communities (Hernandez, 1973; Freedman & Hutchinson, 1976; Chapin & Chapin, 1980; NWT State of the Environment Report, 2022; Malik & Ford, 2025). These regions are also home to 27 First Nations, Métis, and Inuit communities (CIRNAC, 2023); who rely on sustainable land relations and whose land sovereignty has been undermined by past and ongoing resource exploitation (Reyes-García et al., 2019; Kennedy et al., 2023). Many boreal and arctic plant communities, are still recovering from the impacts of resource extraction (Lee & Boutin, 2006; van Rensen et al., 2015). Ecological research in northern habitats poses significant logistic challenges, which emphasises the need for accessible methods to inform effective restoration decisions.

Next generation sequencing (NGS) enables efficient and cost-effective access to comprehensive genetic markers to complement field studies and to support informed restoration and conservation, particularly in northern regions where field studies are less tractable (Ekblom & Galindo, 2011; Breed et al., 2019; Heuertz et al., 2023; Theissinger et al., 2023). In particular, the analysis of single nucleotide polymorphisms (SNP) can help reveal how stochastic and adaptive forces have shaped population differentiation over time and at large geographical scales (Savolainen et al., 2007; Siol et al., 2010; Sork, 2017). Additionally, SNPs can help disentangle taxonomic complexities often observed in widespread non-model organisms (de Villemereuil et al., 2016). To increase sample size, Genotyping-in-Thousands by Sequencing (GT-seq) uses targeted amplicon sequencing to genotype hundreds to thousands of individuals at 50 to 500 SNP loci in a single Illumina Hi-Seq Lane (Campbell et al., 2015). GT-seq has been applied successfully across a wide variety of taxa to address conservation questions including the genetic structure of Western rattlesnakes in Canada (Schmidt et al., 2020), genetic stock identification and kinship analysis of walleyes in the Great Lakes (Euclide et al., 2022), population monitoring using fecal samples of polar bears in the Canadian arctic (Hayward et al., 2022), and the optimization of marker assisted breeding for pepper species in the *Capsicum* genus (Jo et al., 2021). By reducing sequencing costs and increasing throughput, methods like GT-seq could help reveal cryptic species and population structure to inform provenance selection for restoration projects.

Here, we introduce a novel GT-seq SNP panel and analysis pipeline to study wild and cultivated populations of three species of grasses in the *Calamagrostis* genus (*C. canadensis, C. stricta* ssp*. inexpansa,* and *C. purpurascens*) collected across the Northwest Territories of Canada. These species are commonly used in restoration due to their high tolerance for disturbed soils, and significance to indigenous and local communities, but are frequently misidentified due to hybridization, morphological variations, polyploidy and apomixis (Lieffers et al., 1993; Marr et al., 2011; Nygren, 2010). We first evaluate whether the ddRAD-seq and GT-seq data yield similar genetic differentiation patterns among wild populations of *C. canadensis, C. stricta* ssp*. inexpansa,* and *C. purpurascens* from the Northwest Territories. Second, we use the GT-seq panel to investigate potential for hybridization among wild populations of these three species. Third, we characterize geographic variation and population structure among wild populations and evaluate whether cultivars of *C. canadensis* used in restoration projects are representative of wild populations. We discuss how resolving population structure and hybridization potential of *Calamagrostis* species in the Northwest Territories of Canada can inform more effective and sustainable restoration efforts, including provenancing best-practices, and fostering reconciliation with the land and local indigenous communities.

## 2. Materials and Methods

### 2.1 Study system

The genus *Calamagrostis* Adans. (reed grass) comprises ∼ 260 species of perennial rhizomatous grasses in the Poaceae family (Stebbins, 1930), with approximately half of identified species found in high-elevation areas throughout the Americas (Soreng & Greene, 2003). Of the 25 species known to occur in North America, *C. canadensis*, *C. stricta* ssp. *inexpansa*, and *C. purpurascens* have the broadest ranges, extending from the southern U.S. into the Arctic (Marr, Hebda, & Zamluk, 2011). Taxonomic classification of *Calamagrostis* species is complicated by variation in ploidy (diploid to decaploid), apomixis, and hybridization, resulting in subtle phenotypic differences (Nygren, 1949; Greene, 1984; Marr et al., 2011), and high misidentification rates in herbarium records (Marr et al., 2011).

Notwithstanding taxonomic difficulties, *Calamagrostis* is an important genus for restoration, particularly in temperate regions. Rhizomatous growth and tolerance to grazing, flooding, and acidic soils make *Calamagrostis* species valuable for erosion control and soil stabilization (Gilman, 1999). These reed grasses also provide food for large wild mammals and livestock. For example, *Calamagrostis canadensis* is a key diet component of bison and elk in the Northwest Territories (Hogg & Lieffers, 1991; Tesky, 1992). We focus on three common species in northern Canada that are known to colonize abandoned coal mines and oil spill sites (Watson, 1989; Tesky, 1992), and therefore are good candidates for restoration projects: *C. canadensis*, *C. stricta* ssp. *inexpansa*, and *C. purpurascens*.

### 2.2 Sample collection and tissue sampling

Seeds provided by the Aurora Research Institute (ARI) for this study were sampled from 54 wild populations of *Calamagrostis* across 10 degrees of latitude, 23 degrees of longitude and representing seven major geographic regions of the Canadian Northwest Territories (Table 1, Figure 1). Collections were conducted between 2005-2008 under the licence number 13896 on disturbed areas in the Inuvialuit Settlement Region and east of the Dempster between Inuvik and Tsiigehtchic. Seed lines and voucher specimens are maintained by the Western Arctic Research Centre of the ARI in Inuvik, Northwest Territories. Each seed population was identified at the time of collection based on morphological features as summarized in Table 1. In cases where morphological identification was inconclusive, individuals were labeled as ‘*Calamagrostis sp*’. In addition to natural collections, we obtained eight seed mixes of *C. canadensis* from commercial cultivars in the U.S and Canada (Table 1, Figure 1). Since the geographic origins of commercial populations are unknown, we assign geographic location of each commercial garden as the ‘population’ for the purposes of this study.

**Figure 1.**
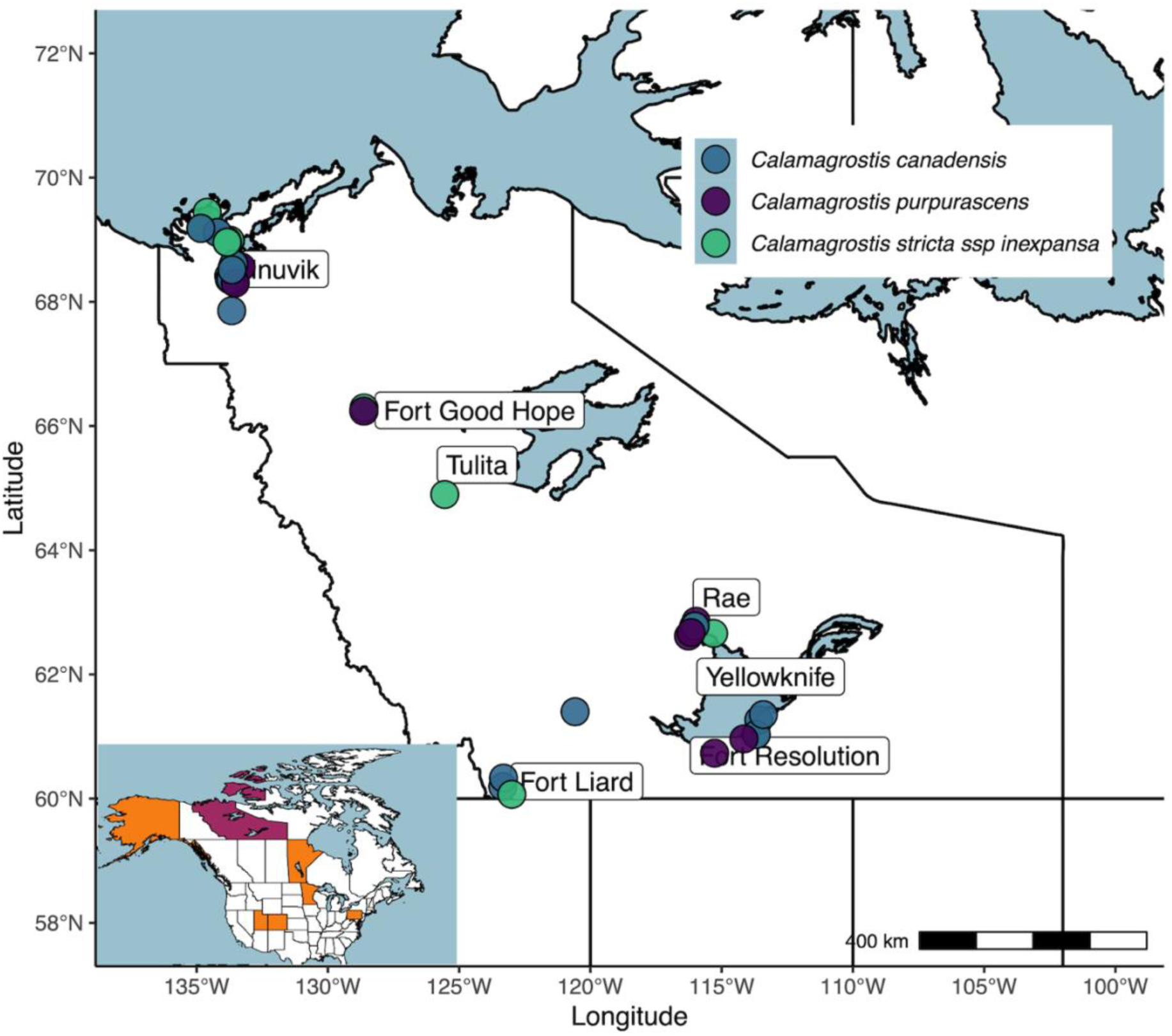
Map of 63 collection sites of three *Calamagrostis* species in the Northwest Territories of Canada. Colour coding for *C. canadensis* (Blue), *C. purpurascens* (Purple) and, *C. stricta* ssp*. inexpansa* (Green) is based on morphological traits. Inset map highlights the Northwest Territories (purple) and the provinces or states from which cultivars were obtained (Orange).

**Table 1.**
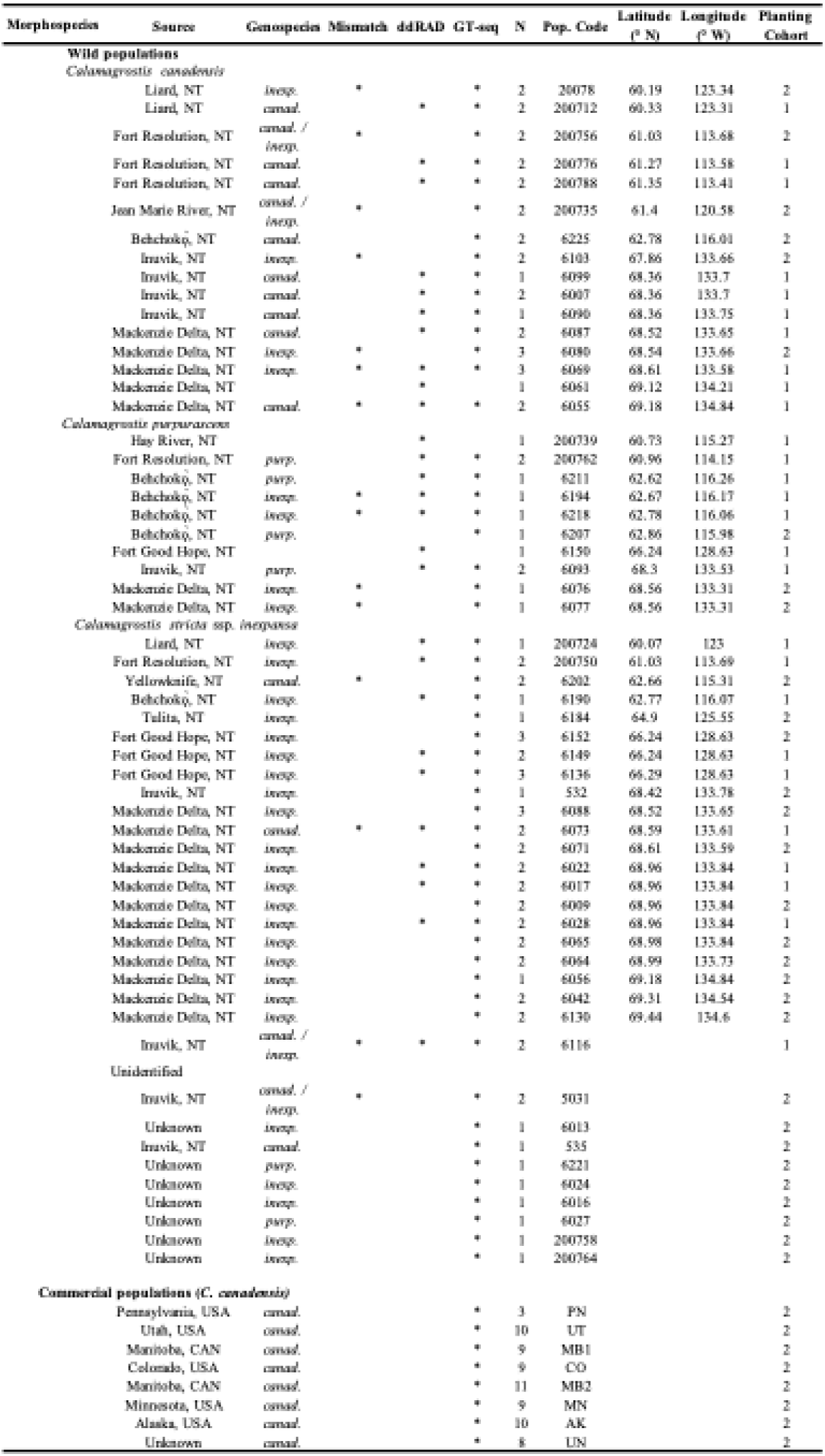
Population summary of three *Calamagrostis* species (*C*. *canadensis*, *C*. *purpurascens*, *C*. *stricta* ssp. *inexpansa*) collected from wild, cultivated, and unknown sources and shared by the Aurora Research Institute (ARI). Up to 11 individuals (N) were assigned a species based on morphological characteristics (Morphospecies) different from Local Fisher Discriminant Analysis (LFDA) of GT-seq data (Genospecies), with disagreements indicated by an asterisk (Mismatch). For commercial accessions, geographical region represents common garden locations as seed sourcing information is unknown. Seeds of ‘unknown’ source are from lines maintained by ARI of undocumented origin.

Both wild and commercial seed accessions were maintained at 4 °C until seeds from each population were propagated at the Queen’s University Phytotron in Kingston, Ontario. After 15 to 30 days, leaf tissue samples from individual were harvested, flash frozen in liquid nitrogen, and stored at - 80°C for DNA extraction. Seed sowing and tissue sampling occurred in two cohorts. The first cohort composed of 112 individuals across 51 wild and five commercial accessions was germinated in March 2018 to create a double digestion restriction associated DNA sequencing (ddRAD-seq) library, and to develop and validate GT-seq primers. The second cohort, composed of 123 individuals across 33 wild and eight commercial accessions, was germinated in March 2022 to apply the GT-seq to more individuals (Table 1).

### 2.3 DNA isolation and library preparation

Genomic DNA was isolated from frozen leaf tissue (0.05-0.1g) using a CTAB protocol (Doyle & Doyle, 1987) modified to include two to three cold prewashes containing CTAB buffer with Polyvinylpyrrolidone PVP but no CTAB lysis agent. This step was added to help remove carbohydrates and other secondary metabolites that often occur at higher concentrations in young leaf tissue and can interfere with extraction chemistry and/or co-precipitate with DNA. The quality and quantity of isolated DNA were assessed using a Denovix DS-11 spectrophotometer. Samples with less than∼10ng/μL were excluded from library preparation.

To construct the ddRAD library from cohort 1, genomic DNA from 11 *C. canadensis*, eight *C. purpurascens*,15 *C. stricta* ssp*. Inexpansa,* four commercial *C. canadensis* and two unknown *Calamagrostis sp* individuals passed quality checks with sufficient DNA for library construction.

Libraries were constructed following published ddRAD protocols (Jensen et al., 2020; Peterson et al., 2012) with *SbfI* (NEB #R3642) and *MluCI* (NEB #R0538) restriction enzymes from New England BioLabs. Briefly, digested fragments were ligated to adaptors corresponding to each enzyme overhang, with a sample-specific 4-6bp molecular barcode compatible with *SbfI* and with degenerate 8bp unique molecular identifier (UMI) barcodes added to the 3’ end of DNA fragments, along with a 6bp plate-specific library barcode. The ligated fragments were pooled and size-selected for a range of 400 to 490bp using a BluePippin 2% agarose cassette with internal marker V1 (Sage science). Following PCR amplification (Jensen et al., 2020), the resulting ddRAD library was sequenced at Génome Québec, Canada in an Illumina HiSeq 4000 partial flow cell with 2 × 150 bp paired end reads at a loading concentration of 360 pM library, producing a total of 210,819,849 raw reads. The GT-seq library preparation for cohort 2 is described in a later section, following description of the GT-seq marker development.

### 2.4 Genotyping and SNP calling

Single nucleotide polymorphisms (SNPs) from the ddRAD data in cohort 1 were identified with the STACKS v.2.3 (Catchen et al., 2011) pipeline running on high-performance computing clusters maintained by BCNet, Compute Canada (now Digital Alliance; https://alliancecan.ca/), and the Center for Advanced Computing at Queen’s University. Demultiplexing, filtering and trimming of reads was performed on STACKS v 2.3 using the *process_radtags* pipeline with the barcode rescue option (-r) and only the *SbfI* enzyme. The cut site of *MluCI* was excluded because it does not appear at the beginning of the second read due to the addition of degenerate inline barcodes. We used the degenerate barcodes on the 3’ end of the DNA library to identified and removed DNA duplicates using the *ParseDBR_ddrad* pipeline (https://github.com/Eljensen/ParseDBR_ddRAD). We used the Python script *ParseFastQ.py* for each FASTA file with the arguments -*-enzyme* = AATT for the second restriction enzyme of MluCI, *--index* = GGCATC corresponding to the 5’ end of the degenerate barcode sequence (i.e., not the Illumina barcode index), and the anchor enzyme sequence length *–hdrlen* = 3. The *de novo* assembly and variant calling were performed using the *denovo_map.pl* pipeline in STACKS v.2.3 using the arguments *M = n* = 5 and *m* = 3. Lastly, the *populations* module in STACKS was run with the parameters *p* = 1 and *r* = 0.2 for an initial filtering of SNPs. These parameters filter for loci present in at least one species, and at least 20% of the individuals within that species group.

Further SNP filtering was performed using *VCFtools* v.1.16 (Danecek et al., 2011) to remove low-confidence genotypes with a minimum Phred quality score of 20, a sequencing depth of at least 5 reads per genotype and a minimum average of 5 reads per site, with a minor allele frequency (MAF) of greater than 0.02. Finally, we removed indels and retained only SNPs for individuals and loci with less than 50% missing data.

### 2.5 ddRAD data analysis

Data analysis for the ddRAD data was performed in *R* version v.4.2.3 with *R Studio* v.2023.06.2 (R Core Team, 2020). To identify and visualize genetic similarity, we converted the VCF file into a *genind* object using the *vcfR* v.1.14 package (Knaus & Grünwald, 2017). We then applied non-supervised machine learning algorithms to construct a Principal Components Analysis (PCA) with package *adegenet* v.2.1.10 (Jombart & Ahmed, 2011). To quantify genetic differentiation among species, we calculated global and pairwise Nei’s *F_ST_* using the package *hierfstat* v.05-11(Goudet, 2005). Using pairwise *F_ST_* values, we visualized genetic distance using a neighbour joining algorithm in the *ape* v.5.7-1 package (Paradis & Schliep, 2019). A main goal of our data analysis was to develop a reproducible and computationally efficient pipeline with minimal program dependencies. Therefore, instead of using the commonly used program *STRUCTURE* (Pritchard et al., 2000) for population structure inference, we instead employed the *LEA* v3.10.2 (Frichot & François, 2015) package in R. This program uses non-negative matrix factorization algorithms that provide outputs similar to those from *STURCTURE* (Frichot & François, 2015). We ran the *snmf* command to estimate individual ancestry and admixture coefficients using five repetitions with 1000 iterations each for each level of *K* (1 to 10) with an alpha regularization parameter of 50. To assess predictive performance of each value of *K*, we evaluated the cross entropy for each run at each level of *K* and selected the run and level with the lowest cross-entropy as the best estimate of *K* (Supplementary Figure 1A). We graphically visualized the Q-values for each individual from the optimal *K* value using *ggplot2* v.3.5.1 (Wickham, 2016).

### 2.6 GT-seq panel development, optimization and genotype calling

To help determine which loci to include in the GT-seq panel, we first used *VCFtools* to calculate summary statistics from the filtered ddRAD SNP data. We removed sites with a minimum sequencing depth below 5 and a mean site depth across individuals less than 7. We also excluded loci with more than 1000 reads to exclude potential organellar loci and high-copy-number paralogues (e.g., pseudogenes). Of the remaining loci, we selected those with less than 70% missing data and a minor allele frequency (MAF) of at least 0.015, resulting in 703 candidate loci to target with GT-seq primers. Sequences for these loci were combined into a single FASTA file and sent to GTseek LLC for custom primer design. Since each locus may contain multiple SNPs, primers were designed to target the SNP with the highest MAF in each target sequence. Of the initial 703 selected SNPs sent to GTseek, 543 included SNPs appropriate for primer design. After removing low-quality primers based on *in silico* analysis (e.g., low binding temperature, multiple binding targets) we retained 347 primer sets for testing.

To validate and optimize the 347 GT-seq primers, we sequenced a pilot library of 45 *Calamagrostis* individuals, including 25 individuals used in the ddRAD library, eight individuals from five commercial populations, and 14 individuals from cohort 1 that were not initially included in the ddRAD library. The dual index library was constructed following Campbell et al. (2015) with GT-seq primers that included individual indices for demultiplexing of samples, and regions needed for sequencing on Illumina sequencers. This pilot library was sequenced using Illumina MiSeq at the Kingston Health Sciences Centre (KHSC) located in Kingston, Ontario Canada. We used the MiSeq Reagent Kit v3with paired-end 2×75 bp at a loading concentration of 8 pM library with 2% PhiX spike-in. SNPs from the primers sets was assessed at GTseek LLC using the *GTseq_SeqTest.pl* and *GTseq_Primer-Interaction-Test.pl* custom scripts available from the GT-seq pipeline (https://github.com/GTseq/GTseq-Pipeline) (Campbell et al., 2015). Of the initial 347 custom locus primers, we removed 18 loci from the final GT-seq panel due to an overabundance of off-target sequences in the dataset containing their primer sequences (i.e., > 1% of total reads).

We applied the final GT-seq panel of 329 loci to a library of genomic DNA isolated from 95 filed-collected individuals (32 *C. canadensis*, 11 *C. purpurascens*, 42 *C. stricta* and 10 unknown *C. sp.*) and 74 commercial cultivar *C. canadensis* samples from cohort 1 and 2. Twenty-five individuals from the ddRAD library from cohort 1 were included in the final GT-seq library to measure consistency between ddRAD and GT-seq (Table 1). We again constructed a sequencing library following standard protocol from Campbell et al. (2015) and sequenced it on an Illumina MiSeq at the KHSC using 2 × 75 bp at a loading concentration of 8 pM library and 2% PhiX.

We modified the standard GT-seq pipeline scripts (https://github.com/GTseq/GTseq-Pipeline) (Campbell et al., 2015) with modifications to account for variation in reported ploidy in the *Calamagrostis* genus (2 × to 10 ×). Specifically, we modified the *GTseq_Genotyper_v3.pl* script to change the allele ratio thresholds for homozygotes from 10:1 to 100:1 to account for differences in binomial sampling probabilities expected lower frequency of minor alleles in polyploids compared to diploids. We also excluded loci with less than 10 reads for the minor allele, to avoid an inflation of heterozygosity that might occur in low-coverage loci given these new thresholds. Individual genotypes were then compiled into a matrix of genotypes for each individual in CSV format using the *GTseq_GenoCompile_v3.pl* script (Campbell et al., 2015).

### 2.7 GT-seq panel validation and data analyses

We applied additional quality controls to the final GT-seq panel by removing fixed loci, loci represented by fewer than 50% of individuals and individuals with fewer than 50% of loci. To compare sequencing methods (ddRAD and GT-seq) and validate our targeted SNPs for species identification, we divided our dataset into three subsets for analysis. The first analysis subset, which we refer to as the ‘validation set’ contained 29 individuals (14 *C. canadensis* from 8 populations, 5 *C. purpurascens* from 3 populations and 10 *C. stricta* ssp*. inexpansa* from 6 populations). This subset included 17 individuals used in the ddRAD analysis, as noted above (Table 1). We refer to the second analysis subset as the ‘wild’ dataset because it includes 93 field-collected individuals genotyped by GT-seq (30 *C. canadensis*, 10 *C. purpurascens,* 42 *C. stricta* ssp*. inexpansa,* and 10 unknown *Calamagrostis sp.*, based on morphological identification), including the 29 individuals in the validation set. The third ‘full’ dataset combines all GT-seq loci from the ‘wild’ dataset with GT-seq loci from 69 individuals sampled from eight commercial populations. Finally, a fourth ‘commercial’ dataset containing only *C. canadensis* individuals from field-collected and commercial populations.

Analysis of the GT-seq markers was performed in R version v.4.3.2 with R Studio v.2023.06.2 (R Core Team, 2020). We first imputed individual missing loci by species using *randomForest* v.4.7 (Liaw & Wiener, 2002) with 10 iterations and 5000 trees and converted each of the four datasets (i.e., validation, wild, full, commercial) into *genind* objects using the *vcfR* v.1.14 (Knaus & Grünwald, 2017) package. To compare the GT-seq and ddRAD data, we performed a Principal Components Analysis (PCA) of the validation dataset using *adegenet* v.2.1.10 (Jombart & Ahmed, 2011) and visualized genetic similarity of PC scores using *ggplot2* v.3.5.1 (Wickham, 2016). To assess genetic differentiation among the three species in the validation dataset, we calculated global and pairwise Nei’s *F_ST_* in the *hierfstat* v.05-11 package (Goudet, 2005), and used these to construct a neighbour joining tree with *ape* v.5.7-1 (Paradis & Schliep, 2019). We also used the *LEA* package v3.10.2 (Frichot & François, 2015) to estimate individual ancestry and admixture coefficients using five replicates of 1000 iterations for each level of *K* (1 through 10) at an alpha regularization parameter of 50; we selected *K* with the lowest cross entropy (Supplementary Figure 1B). We visualized ancestry coefficients with *ggplot2* using the Q-values for each individual from the optimal *K* run.

After comparing the GT-seq with ddRAD results in the validation dataset, we used the GT-seq loci to analyze genetic differentiation and individual clustering in the wild, full and commercial datasets. To visualize genetic variation among individuals, we plotted PC scores from a PCA in *adegenet* v.2.1.10 (Jombart & Ahmed, 2011). To account for potential errors in species classification based on morphology, we conducted a Local Fisher Discriminant Analysis (LFDA) to predict the correct species assignment. An LFDA is a supervised machine learning algorithm commonly used for dimensionality reduction while accounting for the local structure of the data (Sugiyama, 2006). We used the ‘validation’ dataset to train an LFDA model using a *Mabayes* tolerance of 10^-4^, two nearest neighbors, and two reduced features with cross-validation to identify the optimal model. The results of this training LFDA model were input into the *pred.lfda* function from the DA v.1.2 (Tang & Li, 2019) package to predict species identity for each individual in the wild and full datasets. Specifically, the predicted species of each sample was assigned based on posterior probabilities from a Naïve bayes classifier, which assumes conditional independence of the genetic markers. The z-scores of the LFDA model and the posterior probabilities of the predicted species identities were visualized with *ggplot2* (Wickham, 2016). Using the predicted identities from the LFDA model, we re-ran the PCA, LEA ancestry and admixture analyses as described above for the for the wild, full and commercial datasets.

In addition to visualizing genetic variation based on PC scores, we also calculated global and pairwise Nei’s *F_ST_* values and heterozygosity to measure genetic differentiation among individuals. Using the pairwise *F_ST_* values as a measure of genetic distance, we constructed a neighbour joining tree of the wild dataset using *ape* v.5.7-1 (Paradis & Schliep, 2019). Finally, to test for geographic structure among populations within each species in the wild dataset, we conducted an Isolation-by-distance (IBD) analysis. To do this, we used *poppr* v.2.9.6 (Kamvar et al., 2014) to calculate pairwise Nei *F_ST_* as our estimate of genetic distance, and for geographic distance estimates we calculated pairwise Euclidean distances from latitude and longitude coordinates of each individual using *stats* v.4.3.2 (R Core Team, 2020). Finally, we used the pairwise genetic and geographical distance matrices in a Mantel test of the Pearson correlation coefficients as implemented in *vegan* v.2.6-4 (Oksanen et al., 2022). R codes for all of these analyses can be found at (https://github.com/gomezquijano/Calamagrostis_PPLANTs).

## 3. Results

### 3.1 ddRAD data

From the assembly of ddRAD data, we identified a total of 252,161 SNPs across 47,845 loci and 40 individuals. After filtering, this was reduced to 9,152 SNPs across 2,951 loci for 27 individuals (10 *C. canadensis*, 7 *C. purpurascens* and 10 *C. stricta* ssp. *inexpansa)* from cohort 1. Visualizing the first two principal component (PC) scores revealed distinct genetic clusters among *C. canadensis*, *C. stricta* ssp*. inexpansa* and *C. purpurascens* species (Figure 2A). Additionally, we found genotypes consistent with species misidentification and interspecific hybridization (Figure 2A-B). Specifically, we found three individuals that were genetically similar to *C. stricta* ssp*. inexpansa* but were identified as *C. canadensis* (N = 2) or *C. purpurascens* (N = 1) based on morphology (Figure 2A). Additionally, two individuals initially identified in the field as *C. purpurascens* and *C. stricta* ssp*. inexpansa* were genetically intermediate to the three species clusters (Figure 2A).

**Figure 2.**
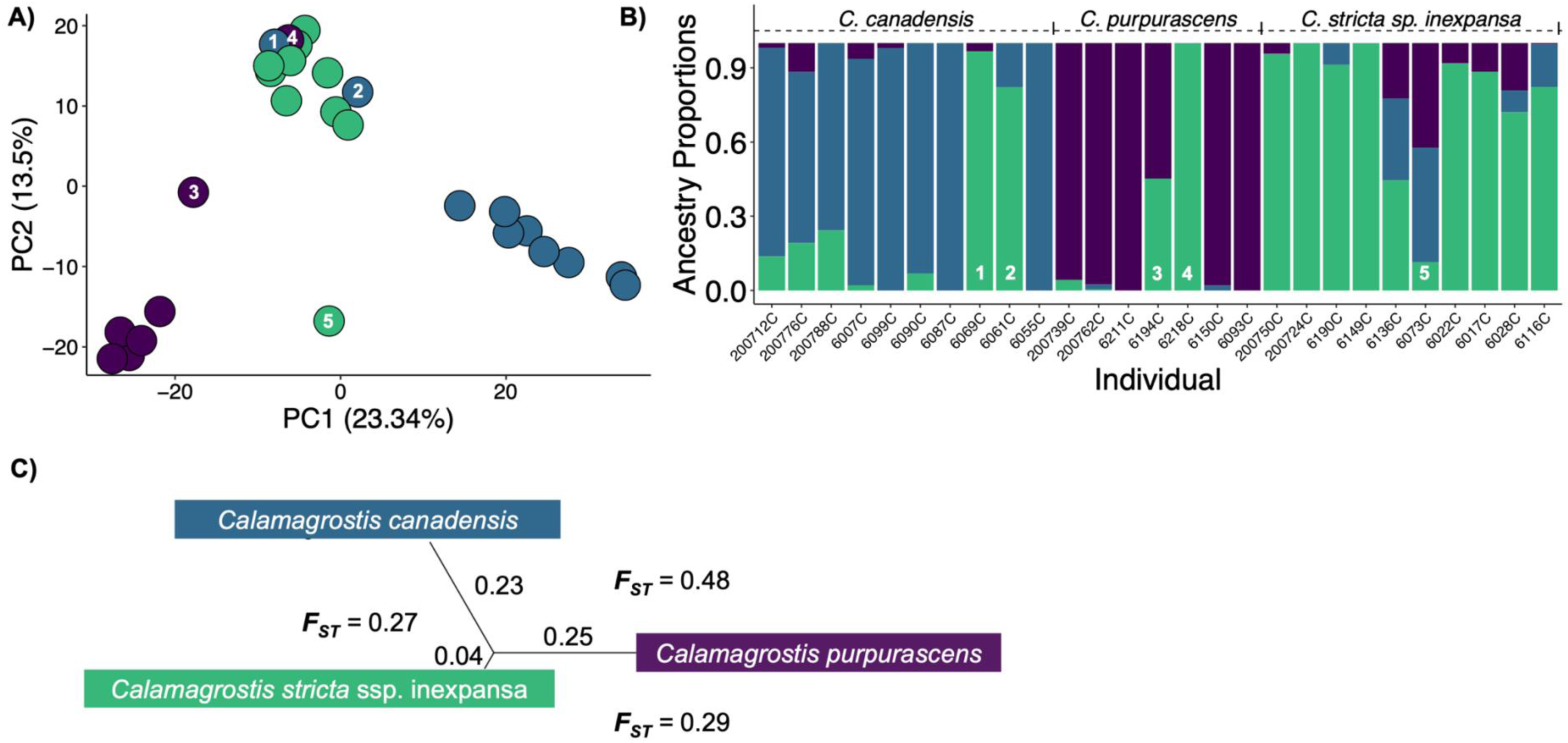
Genetic structure analysis of 9,152 SNPs from ddRAD data for 27 field-collected individuals representing three species of *Calamagrostis*. A) Principal component analysis (PCA) colour coded based on morphological characteristics: *C. canadensis* (Blue), *C. purpurascens* (Purple) and, *C. stricta* ssp*. inexpansa* (Green). Numbered points (1-5) are individuals whose morphological species identification did not match other individuals in the same principal component cluster. B) Ancestry proportion plot implemented in the LEA package at *K=*3 and arranged from southern to northern latitude within a species. Numbered individuals (1-5) show ancestry proportions of misidentified and miss-clustered individuals in panel A. C) Neighbour-joining tree and pairwise *F_ST_* statistics for each species. Branch length and adjacent values represent genetic distances between each grouping. *F_ST_* values are shown for each species pair.

An analysis of ancestry and admixture using LEA supported three genetic clusters (optimal *K =* 3, Supplementary Figure 1A) with each group corresponding to one of the three study species (Figure 2B). The misidentified and hybrid genotypes identified in the PCA were also evident in the LEA results. Specifically, the three misidentified individuals in the PCA had *C. stricta* ssp*. inexpansa* ancestry coefficients > 80% (Figure 2B). One individual identified as *C. purpurascens* and genetically intermediate to *C. purpurascens* and *C. stricta* ssp. *inexpansa* in the PCA also had 50% ancestry from *C. stricta* ssp*. inexpansa* and 50% from *C. purpurascens* in the LEA results, consistent with a first-generation hybrid individual. Another individual that was identified as *C. stricta* ssp. *inexpansa* but genetically intermediate to *C. purpurascens* and *C. canadensis* in the PCA similarly showed 50% *C*. *canadensis* and 50% *C. purpurascens* ancestry in the LEA, also consistent with an F1 hybrid (Figure 2B). In addition to the likely F1 hybrids, the LEA analysis revealed moderate levels (20-30%) of mixed ancestry in several individuals, consistent with introgression from *C. stricta* ssp. *inexpansa* into southern *C. canadensis* individuals, and lower levels (10-20%) of introgression from *C. canadensis* and *C. purpurascens* into Northern *C. stricta* ssp*. inexpansa* individuals (Figure 2B).

Consistent with the LEA results, genetic differentiation metrics identified *C. purpurascens* and *C. canadensis* as the most genetically distinct among the three species. Global *F_ST_* among all species was 0.35, with interspecific pairwise *F_ST_* ranging from 0.27 between *C. canadensis* and *C. stricta* ssp. *inexpansa*, and 0.29 between C*. stricta* ssp*. inexpansa* and *C. purpurascens* up to 0.48 between *C. canadensis* and *C. purpurascens* (Figure 2C). Genetic distances from the neighbour joining tree ranged from 0.04 for *C. stricta* ssp. *inexpansa*, to 0.23 for *C. canadensis* and 0.25 for *C. purpurascens* (Figure 2C).

### 3.2 GT-seq panel validation

The initial GT-seq SNP panel included 329 loci. After SNP quality filtering of the validation data set, we retained 256 loci across 29 individuals from cohort 1 (for details on individuals see Methods 2.7 and Table 1). This filtering included the removal of misidentified and hybrid individuals identified in the ddRAD data (Figure 2). Principal component analysis of the GT-seq validation dataset revealed three distinct clusters corresponding to the three study species (Figure 3A), and these same clusters were recovered in the LFDA of the three species (Figure 3B). The LEA ancestry analysis revealed similar clustering with the lowest cross entropy at *K =* 3 (Supplementary Figure 1B) corresponding to the three species (Figure 3C). Similar to the ddRAD results (Figure 2B), we also observed moderate (15-35%) *C. stricta* ssp. *inexpansa* ancestry in southern individuals of *C. canadensis*, but less evidence for admixture in *C. purpurascens* (Figure 3C). *Calamagrostis stricta* ssp*. inexpansa* also had low (∼10%) *C. purpurascens* ancestry and moderate (10-25%) *C. canadensis* ancestry in the GT-seq data (Figure 3C).

**Figure 3.**
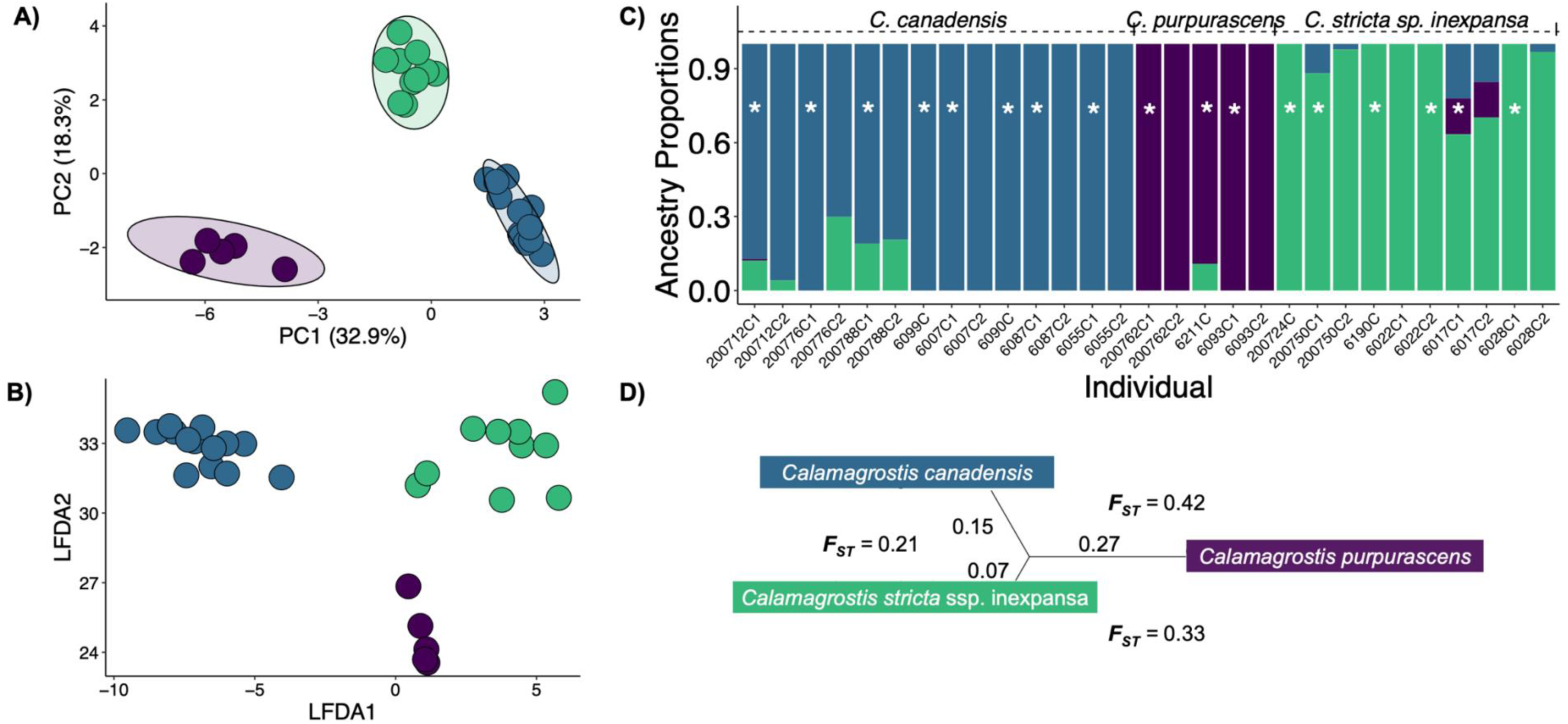
Genetic structure analysis of 256 SNP markers from GT-seq data for the validation dataset of 29 field-collected individuals of three species of *Calamagrostis*. A) PCA with 95% CI ellipses of field-collected *C. canadensis* (Blue), *C. purpurascens* (Purple) and, *C. stricta ssp. inexpansa* (Green) individuals. B) Local Fischer Discriminant analysis (LFDA) clustering of the individuals described in panel (A) used as a training dataset for species identification. C) Ancestry proportion plots with *K* = 3 are arranged from southern to northern latitude within a species. Asterisk denotes one of 18 individuals representing individuals from the ddRAD analyses, which are included with technical replicates to test the accuracy and repeatability of species assignment using GT-seq loci. D) Neighbour joining tree and pairwise *Fst* statistics for each species. Branch length and adjacent values represent genetic distances between each grouping. *Fst* values are shown for each species pair.

Global *F_ST_* among species was 0.31 for the GT-seq data, similar to the 0.35 value from the ddRAD data. Interspecific pairwise *F_ST_* was also similar, with 0.21 between *C. canadensis* and *C. stricta* ssp. *inexpansa*, 0.33 between C*. stricta* ssp*. inexpansa* and *C. purpurascens* and 0.42 between *C. canadensis* and *C. purpurascens* (Figure 3D). Lastly, the genetic distances based the neighbour joining tree were 0.07 for *C. stricta ssp. inexpansa*, 0.15 for *C. canadensis* and 0.27 for *C. purpurascens* (Figure 3D), reinforcing higher differentiation between *C. purpurascens* and the other species. Overall, the ddRAD and GT-seq datasets revealed very similar genetic structure (compare Figures 2 & 3).

### 3.3 Species differentiation and hybridization of wild genotypes

After filtering, the GT-seq panel of the wild dataset included 256 loci across 93 field-collected individuals from cohort 1 and 2, which were morphologically identified as 30 *C. canadensis*, 10 *C. purpurascens,* 42 *C. stricta* ssp*. inexpansa,* and 10 unknown *Calamagrostis* species. The commercial dataset included 69 *C. canadensis* individuals from 8 different commercial cultivars on top of the field-collected *C. canadensis* individuals, resulting in a full dataset of 162 individuals. The PCA of the wild dataset revealed the same three distinct species clusters identified in the ddRAD and GT-seq validation datasets (Figure 4A). In contrast, the clusters observed for *C*. *canadensis* and *C*. *stricta* ssp. *inexpansa* along the first to PC axes were less distinct in the GT-seq dataset, with several misidentified (29 of 93) individuals and potential hybrid (19 of 93) individuals of intermediate genotypes between the main *C. Canadensis* and *C. stricta ssp. inexpansa* clusters (Figure 4A).

**Figure 4.**
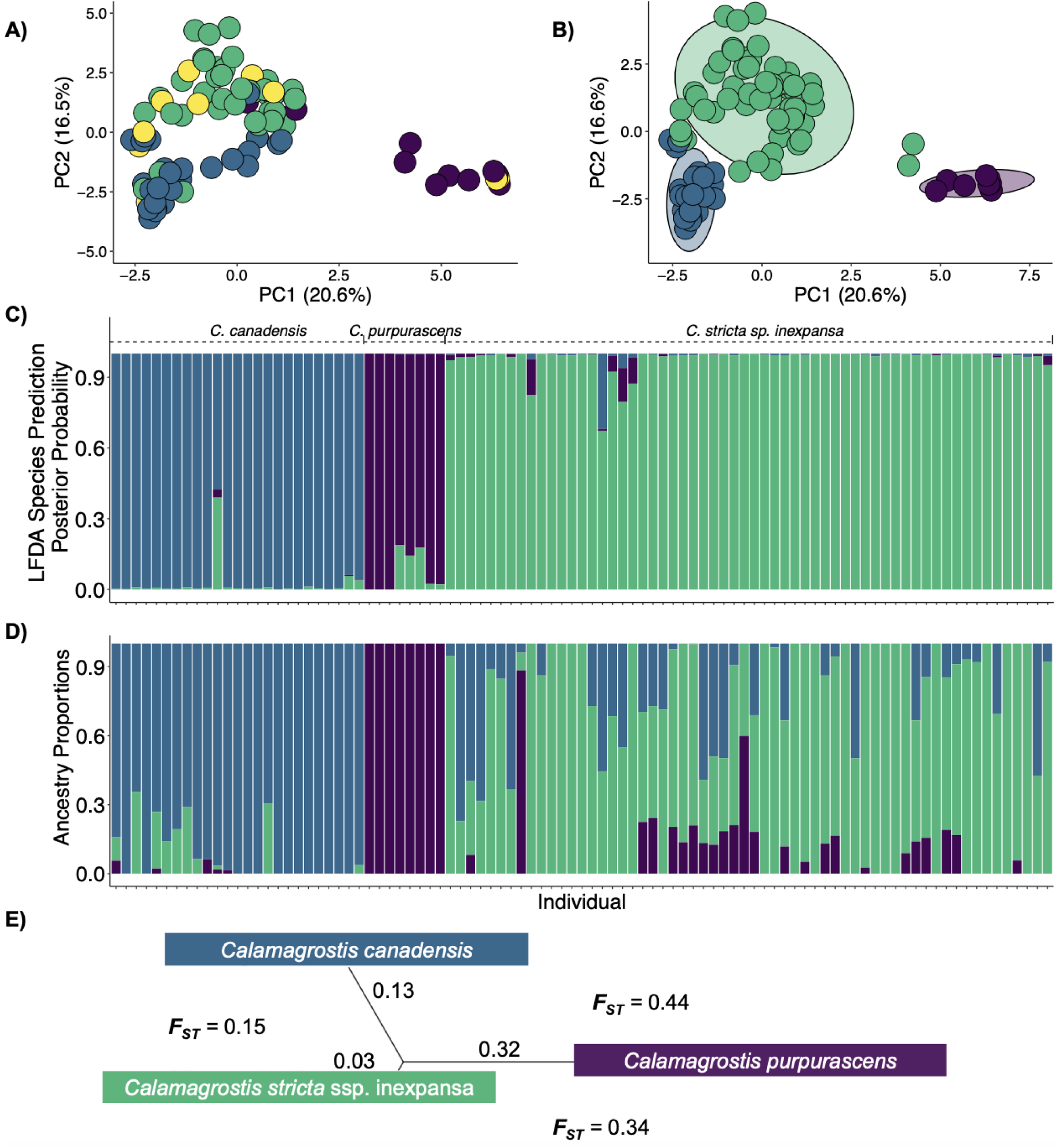
Genetic differences based on 256 SNP markers from GT-seq data for the wild dataset of 93 field-collected individuals of three *Calamagrostis species*. A) Principal component analysis (PCA) of individuals colour-coded by morphology-based species identification: *C. canadensis* (Blue), *C. purpurascens* (Purple), *C. stricta* ssp*. inexpansa* (Green) and unassigned or uncertain *Calamagrostis* species (Yellow). B). PCA with 95% CI ellipses using the LFDA predicted species assignment. C) Posterior probabilities of species assignment based on Naive Bayes Classifier for each individual in the LFDA model. D) Ancestry proportion plot at *K=*3 arranged from southern to northern latitude and LFDA species prediction. E) Neighbour joining tree with branch length and adjacent values represent genetic distances between species. *F_ST_* values are shown for each species pair.

Using the validation dataset as a training model we performed species identity prediction for our wild dataset using an LFDA. The LFDA model predicted 25 *C. canadensis*, 8 *C. purpurascens,* 60 *C. stricta* ssp*. inexpansa* individuals (Figure 4B). Because our LFDA model is a supervised machine learning algorithm it can only predict species identities based on the three categories provided (*C. canadensis, C. stricta* ssp*. inexpansa* and *C. purpurascens*) and therefore potential hybrids were not detected. Instead, we include confidence interval (CI) ellipses in our PCA plot to flag potential hybrids (Figure 4B). Specifically, we identified two individuals that were classified as *C. stricta* ssp*. inexpansa* by LFDA but were outside the confidence CI ellipses of the *C. purpurascens* and *C. stricta* ssp*. inexpansa* clusters (Figure 4B). Similarly, six individuals identified by LFDA as *C. canadensis* and seven identified as *C. stricta* ssp*. inexpansa* fell outside the CI ellipsis. Lastly, the three *C. stricta ssp. inexpansa* individuals with the highest PC2 score fell on the edge of the CI ellipse but not in between any clusters (Figure 4B).

We examined posterior probabilities from the LFDA Naïve Bayes classifier (Figure 4C) and the LEA ancestry coefficient analysis at *K = 3* (Figure 4D) to evaluate the accuracy of LFDA species predictions and identify potential hybrids. The posterior probabilities of the model’s Naïve Bayes classifier approach in the LFDA were > 0.8 for every individual, with two exceptions. The first was classified as a *C. canadensis* individual with (∼ 0.4) probability of being *C. stricta* ssp. *inexpansa*, and the second one was classified as a *C. stricta* ssp*. inexpansa* individual with (∼ 0.3) probability of being *C. canadensis* (Figure 4C). The latter also had high levels of admixture with *C. canadensis* (∼50%) in the LEA ancestry analysis, but the *C. canadensis* individual did not have significant co-ancestry with *C. stricta ssp. inexpansa*. The ancestry analysis also revealed that two individuals predicted as *C. stricta* ssp. *inexpansa* but falling in between the *C. stricta* ssp. *inexpansa* and *C. purpurascens* clusters had > 50% ancestry of *C. purpurascens* (Figure 4BD). Finally, the 10 predicted *C. stricta* ssp*. Inexpansa* individuals falling outside of the CI ellipses in the PCA analyses were found to have > 40% *C. canadensis* ancestry, while the six *C. canadensis* individuals falling outside of the CI ellipsis had > 25% *C. stricta* ssp. inexpansa ancestry (Figure 4D).

The global *F_ST_* among all individuals in the wild dataset was 0.23, with interspecific pairwise *F_ST_* of 0.15 between *C. canadensis* and *C. stricta* ssp. *inexpansa*, 0.34 between C*. stricta* ssp*. inexpansa* and *C. purpurascens* and 0.44 between *C. canadensis* and *C. purpurascens* (Figure 4E). Genetic distances from the neighbour joining tree analysis were 0.03 for *C. stricta* ssp*. inexpansa*, 0.13 for *C. canadensis* and 0.32 for *C. purpurascens* (Figure 4E). Lastly, species-specific mantel test results showed significant isolation-by-distance for *C. canadensis* (r^2^ = 0.23, p-value = 0.002) and *C. stricta* ssp*. inexpansa* (r^2^ = 0.16, p-value = 0.039) but not for *C. purpurascens* (r^2^ = 0.34, p-value = 0.099) (Figure 5).

**Figure 5.**
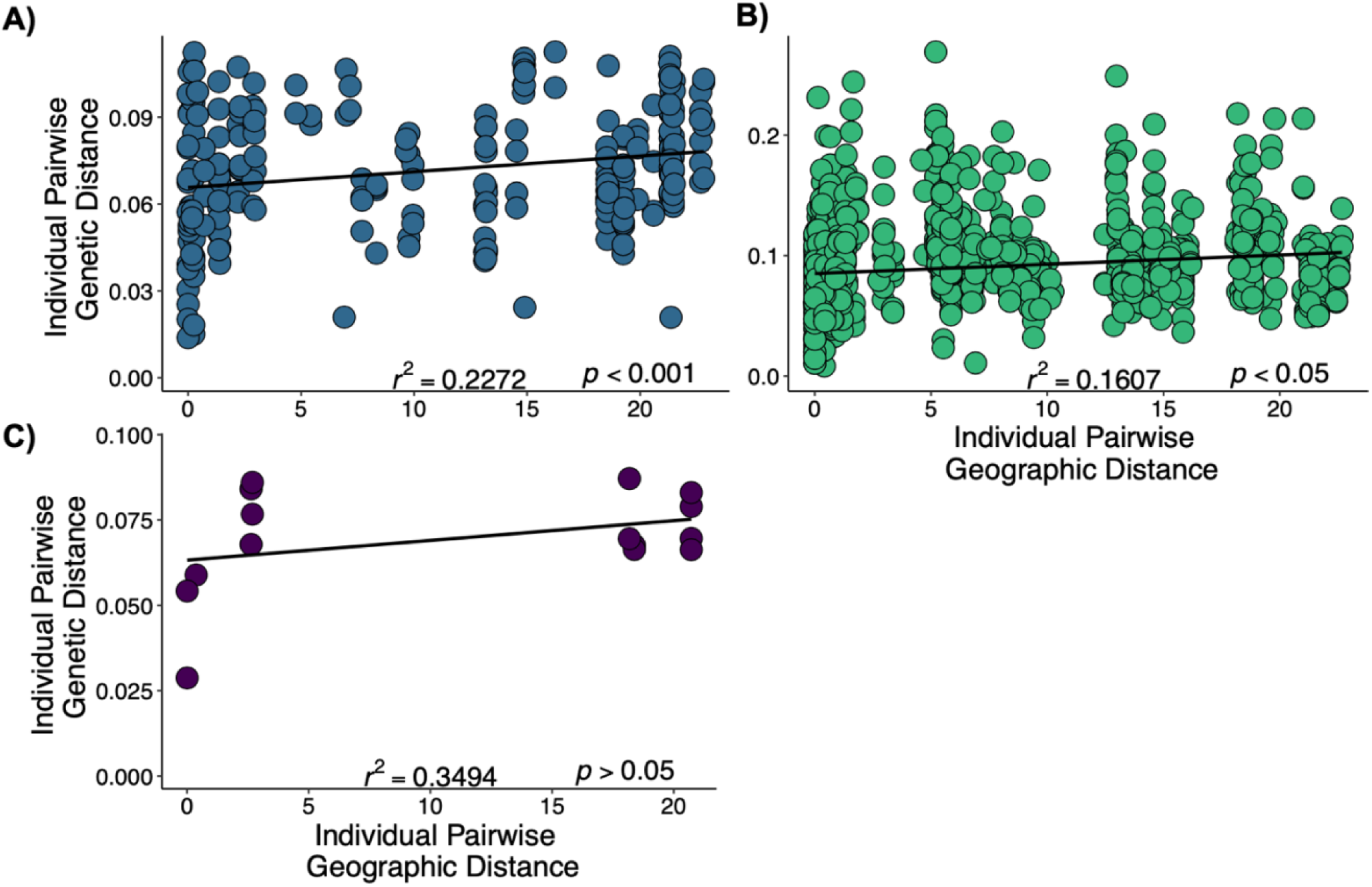
Isolation by distance (IBD) analysis using the wild dataset of species :A) *C. canadensis* (Blue), B) *C. purpurascens* (Purple) and C) *C. stricta* ssp*. inexpansa* (Green) assigned by LFDA predictions rather than morphological characteristics. Each color point for each panel represents a pair of individuals, with Nei’s genetic distance plotted against the individual pairwise geographic distance (latitude × longitude Euclidean distance in decimal degrees). Regression line (black) in each panel was performed using a simple linear model method to show the relationship between the two distance measures, with significance shown for Mantel tests (*r*^2^ and *p-value*) in the bottom right for each panel.

### 3.4 Population structure of commercial genotypes

The full dataset included 256 GT-seq loci across 162 individuals, including 93 individuals grown in cohort 1 and 2 from field-collected seed and 69 individuals sourced from eight different commercial cultivars from cohort 2. The LFDA model predicted that the full dataset was composed of 24 field-collected and 66 commercial *C. canadensis*, 7 field-collected and 1 commercial *C. purpurascens,* and 62 field-collected and 2 commercial *C. stricta* ssp*. inexpansa* individuals (Figure 6A). Lastly, the commercial dataset included the same 256 GT-seq loci across the 69 individuals sourced from eight different commercial cultivars and 24 field-collected individuals that our LFDA classified as *C*. *canadensis* from the full dataset. The PCA and LEA analyses at *K =* 2 showed two clusters of mixed commercial and field-collected individuals, with southern populations being genetically similar to the cultivars PA, MB2 and MN. Northern populations formed a separate cluster that was genetically more similar to the cultivars UT, MB1, AK, UN and CO (Figure 6BC). The commercial individual from MB2 predicted by the LFDA as *C. stricta* ssp*. inexpansa,* clustered with the other MB2 and southern field-collected individuals. Similarly, the commercial individuals from MB1 and UT, predicted as *C. purpurascens* and *C. stricta* ssp*. inexpansa* respectively clustered with the other MB1, UT and northern field-collected individuals (Figure 6C).

**Figure 6.**
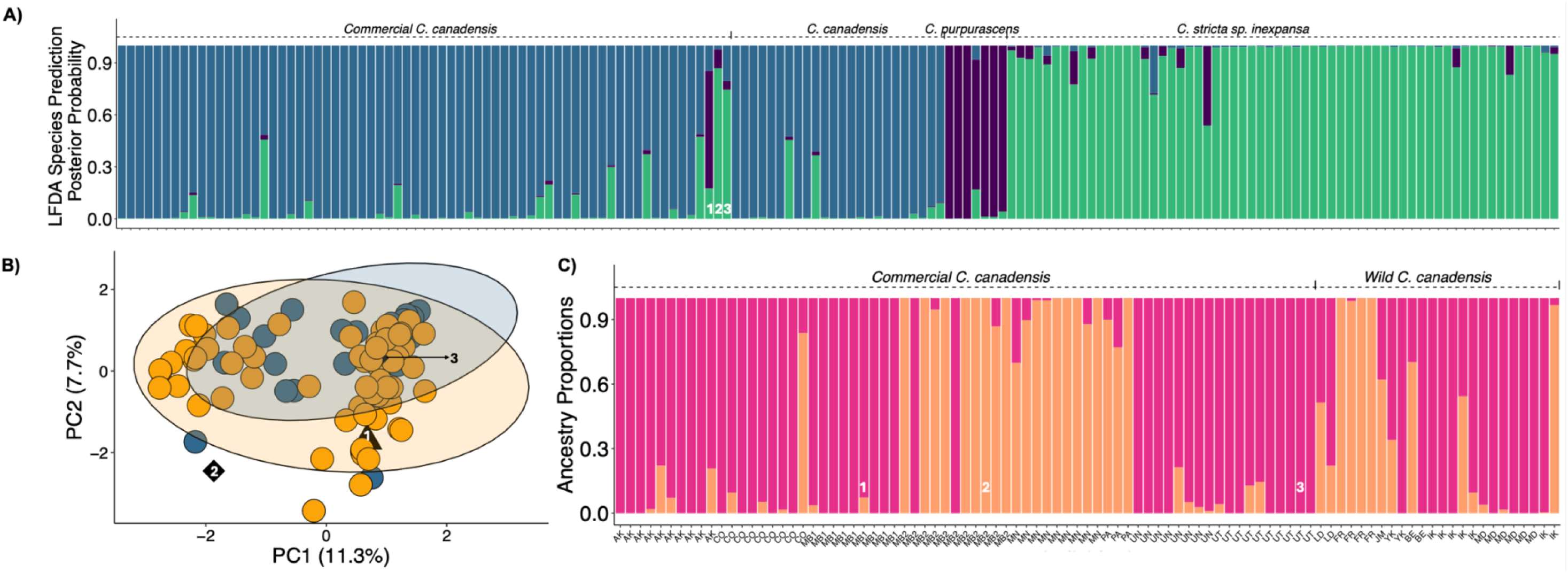
Population and species differentiation analyses using 256 SNP markers from GT-seq data for the full (A) and the commercial (B and C) datasets. A) Posterior probabilities of species assignment based on Naive Bayes Classifier for each individual in the LFDA model using the full dataset. B) Principal component analysis (PCA) of the 69 commercial individuals (Dark yellow) from 8 cultivars and 24 field-collected (Blue) and LFDA predicted *C. canadensis* individuals. The black triangle labeled 1, represent the MB1 individuals predicted as *C. purpurascens* by the LFDA, while the two black diamonds labeled 2 and 3 represent the MB2 and UT individuals predicted as *C. stricta* ssp*. inexpansa* by the LFDA. C) Ancestry proportion at *K=*2 for the commercial dataset. Commercial individuals are arranged by cultivar and labeled by the states or provinces from which they were sourced. Wild individuals are arranged by latitude of origin and labeled as the major geographical region they belong to. Latitude of origin is not recorded for the last two field-collected individuals.

The *F_ST_* between field-collected and commercial *C. canadensis* sources was 0.021, while the *F_ST_* among all commercial individuals was 0.051. Among field-collected populations, heterozygosity ranged from 0.142 to 0.202, with pairwise *F_ST_* values ranging from 0 to 0.231. The heterozygosity for the commercial cultivar populations ranged from 0.115 to 0.172, while the pairwise *F_ST_* ranged from 0 to 0.124. Lastly, the pairwise *F_ST_* between field-collected and commercial cultivars ranged from 0 to 0.234 (Table 2).

**Table 2.**
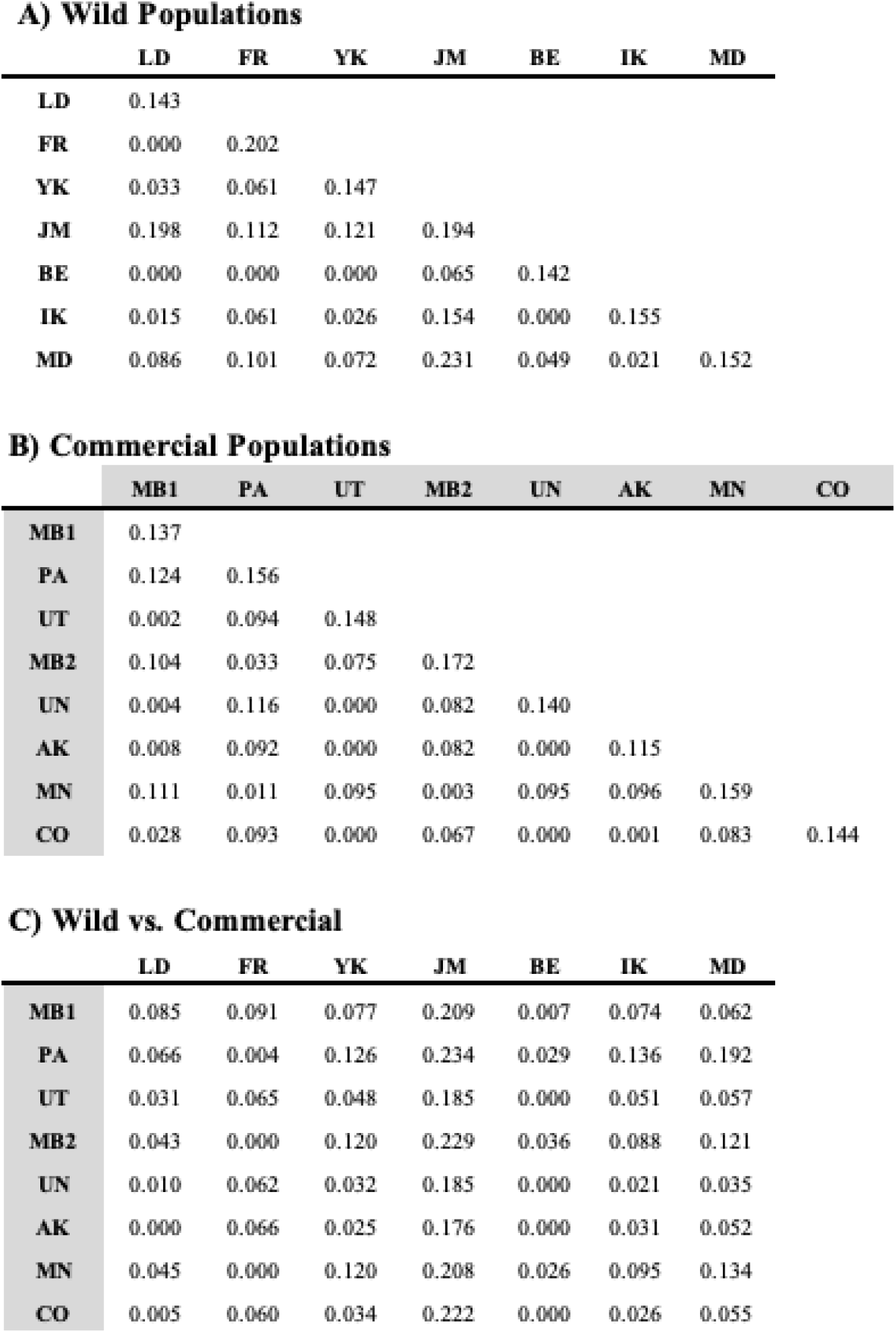
Average observed heterozygosity (diagonal) and pairwise F_ST_ (off-diagonal) from GT-seq data of field-collected (white cells) and commercially cultivated (grey cells) populations of *Calamagrostis canadensis* Wild populations, as listed in Table 1, are grouped by region to increase sample sizes for F_ST_ comparisons (LD – Fort Laird, FR – Fort Resolution, YK – Yellowknife, JM – Jean Marie River, BE – Behchokǫ (formerly Rae-Edzo), IK – Inuvik, MD –Mackenzie Delta). All wild populations are located in the Northwest Territories of Canada.

## 4. Discussion

There is growing demand for habitat reclamation and restoration to help offset the environmental impacts of anthropogenic activity. For example, the United Nations has designated 2021-2030 the “decade of ecosystem restoration” in effort to restore natural landscapes and ecosystems (UNEP, 2021). Establishing clear policies and guidelines for these processes is essential for maintaining healthy ecosystems (Bradshaw, 1984; Palmer & Stewart, 2020). Nonetheless, a major challenge in the field of ecological restoration is determining the best provenancing practices to improve the rate of plant establishment, long-term plant persistence and meet restoration demands (Breed et al., 2013, 2018, 2019; Pedrini et al., 2020). To help address this challenge, multiple provenancing approaches that consider geography (Kiehl et al., 2010, 2014), future climate (Sgrò et al., 2011; Prober et al., 2015), and gene flow (Broadhurst et al., 2008; Breed et al., 2013; Bucharova et al., 2019) when selecting seed sources. Despite high demand for native seed stock, a dearth of genetic data in ecologically important species hinders best practices for provenance sourcing and mixing, especially in northern regions where field trials are limited due to logistic constraints.

To inform restoration in the Northwest Territories of Canda, we developed a targeted SNP panel (N > 250 loci) to confirm morphology-based taxonomic identification, assess genetic differentiation among populations, and investigate potential for hybridization among three species in the *Calamagrostis* genus (*C. canadensis*, *C. stricta* ssp*. inexpansa* and *C. purpurascens*), which are ecologically important and widespread across the Northwest Territories of Canada. Our results show that while morphologically distinct species generally form distinct genetic clusters, some likely have been misidentified, and several intermediate genotypes are consistent with the historical difficulties of morphological identification and recent hybridization respectively. After accounting for species differences and potential hybrid genotypes, we detected isolation-by-distance patterns that can inform provenance strategies for restoration. Given the potential use of *Calamagrostis* species for ecological restoration of eroded and contaminated soils (Tesky, 1992), we also investigated patterns of genetic differentiation between field-collected and commercial cultivar populations of *C. canadensis*. We found that some commercial cultivars were genetically similar to southern wild populations while others were genetically similar to northern wild populations, emphasizing the importance of genetic differentiation data for restoration practices. Below, we discuss the practical applications of GT-seq with machine learning algorithms in non-model systems with complex genomes like *Calamagrostis*. We then examine evidence for historic and contemporary gene flow, population differentiation, and hybridization to inform ecological restoration.

### 4.1 ddRAD-Seq and GT-seq characterize wild populations of polyploid *Calamagrostis* species

Genotyping-in-Thousands by Sequencing (GT-seq) has emerged as a cost-effective and minimally invasive genotyping tool to aid in the conservation and management of non-model systems (Campbell et al., 2015; Euclide et al., 2022; Hayward et al., 2022; Schmidt et al., 2020). However, its accuracy depends on careful panel optimization, previous knowledge of the target species, and clearly defined research questions (Hayward et al., 2022). Non-model systems like *Calamagrostis* and other plants often have large, complex, polyploid genomes, which introduces analytical challenges for GT-seq, ddRAD, and other forms of reduced representation sequencing. In particular, error correction algorithms for SNP calling will typically use minor allele frequency (MAF) with stringent cutoffs that are likely to under-estimate heterozygosity for polyploid loci with polysomic inheritance. For example, consider a diploid heterozygous SNP locus that has a coverage of 100 reads that are sampled following a Bernoulli distribution; the odds of observing MAF less than 10% is quite small under this model (p < 10^-16^). In contrast, the expected minor allele frequency for an octoploid may be as low as 12.5% (i.e., 1/8) and the probability of observing a minor allele frequency of 10% in 100 reads is much higher (p < 0.19). To overcome these challenges we followed similar approaches to those reported for polyploid fish species like Sturgeon (8 × – 12 ×) (Delomas et al., 2021) and Chinook Salmon (3 ×) (Delomas, 2019). By Modifying the allelic ratio threshold to account for polyploidy, we improved accuracy of SNP and multi-locus genotype calling across species while maintaining GT-seq methodology consistent.

A primary goal of GT-seq is to capture relevant genome-wide variation with fewer loci, so that more samples can be included in each sequencing flow cell to reduce costs. Our GT-seq panel of 256 loci included less than 10% of the loci from ddRAD-seq (N = 2,951 loci), yet both retained sufficient genetic variation to resolve species differences (Figures 2 to 4), identify patterns of admixture (Figure 4) and characterize population structure between commercial and field collected *C. canadensis* populations (Figure 6). Therefore, for applications in taxonomy and provenance sourcing for habitat restoration, our panel of GT-seq loci are informative at 1/10^th^ of the sequencing depth and without the need for restriction enzymes.

### 4.2 Evidence for hybridization and population structure

Species ambiguity and misidentification are persistent in the *Calamagrostis* genus, with high misclassification rates in herbarium records dating back to the 1930s (Marr et al., 2011). In particular, morphometric analyses of specimens from British Columbia and adjacent regions have shown that *C. purpurascens* forms a distinct morphological cluster, whereas *C. canadensis* and *C. stricta* frequently overlap, making morphological differentiation difficult (Marr et al., 2011). Consistent with these reported difficulties, ∼1/3 (29 of 93) of specimens in our study were either unidentified (10 of 30) or were morphologically assigned to a species that did not match its multilocus genotype (19 of 30) (Table 1, Figure 4A). Also consistent with the morphological findings of Mar et al. (2011), genetic differentiation was less distinct for *Calamagrostis canadensis* and *C. stricta* ssp. *inexpansa*, compared to *C. purpurascens* which formed a more genetically distinct cluster (Figures 3, 4). However, we note that species form distinct genetic clusters (Figure 2) with less overlap when compared to morphological features (compare to Figure 1 in Mar et al., 2011), indicating potential for molecular markers to assist with taxonomic identification.

To help address challenges with morphology-based species identification, we used an LFDA to predict species identity from GT-seq SNPs with 96% accuracy (Figure 4, Figure 6). Interestingly, individuals with higher levels of admixture were consistently assigned to *C. stricta* ssp. *inexpansa* even when the majority of their ancestry proportion derived from other species (compare Figure 4C vs 4D). This likely reflects bias in the training data, where *C. stricta* individuals had the highest levels of admixture, leading the algorithm to classify admixed individuals as *C*. *stricta*. Resolving hybrids and other admixed individuals will likely require a larger training dataset with more loci, but our results demonstrate that GT-seq markers are sufficient for resolving species and could be particularly useful for identifying seeds and non-reproductive plants with few distinguishing morphological characteristics.

Our ancestry coefficient results diverged from phylogenetic expectations suggesting ongoing and historical gene flow between *C. stricta* ssp. *inexpansa* and *C. purpurascens*. To our knowledge, this has not been observed in previous studies using plastid and nuclear ribosomal (ITS) phylogenetic reconstructions (Peterson et al., 2022; Saarela et al., 2017). Phylogeny and biogeographical analyses have shown that *C. canadensis* and *C. stricta* ssp. *inexpansa* are closely related sister taxa within the same clade (Eurasian), whereas *C. purpurascens* belongs to a distinct lineage (Peterson et al., 2022; Saarela et al., 2017). While our findings largely support this structure, with *C. purpurascens* and *C. canadensis* exhibiting the greatest genetic differentiation and *C. canadensis* - *C. stricta* ssp. *inexpansa* having lower genetic differentiation (Figure 2), we also identified lower than expected genetic differentiation between *C. purpurascens* and *C. stricta* ssp. *inexpansa* regardless of sequencing method. 4D).

Hybridization has been recognized as a major factor in the evolution of *Calamagrostis* species, with evidence for post-speciation gene flow documented among sympatric species in Europe, Asia and North America (e.g., Paszko & Nobis, 2011; Tateoka, 1977; Nygren, 1951). Contemporary gene flow between *C. stricta* and other sister taxa like *C. canesens* has also been reported (Crackles, 1997), and several North American species (e.g., *C. perplexa* and *C. rubescens)* are believed to have hybrid origins involving *C. canadensis*, *C. stricta* and other species during the genus’s circumboreal and transatlantic expansion (Peterson et al., 2022). Consistent with studies of other regions, our analysis of genome-wide SNPs of species in the Northwest Territories of Canada reveal both historical and contemporary introgression between *C. stricta* ssp*. inexpansa* and *C. canadensis*, and between *C. stricta* ssp*. inexpansa* and *C. purpurascens* (Figure 2, Figure 4). Moreover, gene flow between *C. canadensis* and *C. stricta* ssp*. inexpansa* appears to be symmetrical with 15 individuals averaging ∼30% admixed ancestry and seven individuals exhibiting ∼50% ancestry from both species (Figure 4D).

While *C. purpurascens* hybrids have not been widely documented, it is hypothesized that the species originated from complex hybridization events between distinct *Calamagrostis* lineages (Peterson et al., 2022; Saarela et al., 2017). Our analyses revealed one individual with ∼60% *C. purpurascens* and 40% *C. stricta* ssp*. inexpansa* and seven *C. stricta* ssp*. inexpansa* individuals with > 20% ancestry from *C. purpurascens* supporting contemporary hybridization in sympatry (Figure 4D). Similarly, five *C*. *canadensis* individuals showed > 15% ancestry from *C. purpurascens*, suggesting recent gene flow and potential for *C. purpurascens* and *C. canadensis* to hybridize. To our knowledge, this is the first evidence of interspecific hybridization at the genome level involving these three species within the Canadian Arctic and in North America more generally.

Despite evidence for contemporary hybridization among species, we found that genome-wide variation was geographically structured within *C. canadensis* and *C. stricta* ssp*. inexpansa,* but not for *C. purpurascens.* Isolation-by-distance (IBD) is common in wind-pollinated grass species with large geographic ranges, such as *Stipa capillata* in central Europe (Hensen et al., 2010), *Elymus athericus* in European salt marshes (Bockelmann et al., 2003), *Arctophila fulva* in Finland (Kreivi et al., 2005) and *Arrhenatherum elatius* in Germany (Durka et al., 2017). While IBD has not been investigated previously in *Calamagrostis* species in Canada, our sampling spanned a ten-degree latitudinal range across boreal and Arctic ecosystems (Figure 1) allowing us to compare geographically distant populations representing a broad range of environmental characteristics. Therefore, it may not be surprising that we found significant associations between genetic distance and geographic distance (Figure 5). This is particularly evident in *C. canadensis,* where southern populations also exhibited higher admixed ancestry than those from northern regions (Figure 4D, Figure 5A). On the other hand, *C. purpurascens* populations were not significantly differentiated across ∼20 latitude × longitude decimal degrees (Figure 5C), suggesting higher rates of historical and/or contemporary gene flow.

### 4.3 Genetic diversity in cultivated and wild populations

Genetic variation and local adaptation are important factors in successful ecological restoration, particularly for species with large geographical ranges. While we observed overall low genetic differentiation between commercial and field-collected sources (*F_ST_* = 0.021), we found evidence of geographically structured genetic differentiation. Our results identified two genetic clusters within *C. canadensis* that included both cultivar and field-collected populations (Figure 6B,C). The first cluster included three commercial cultivars (MB2, MN, and PA) along with individuals from six wild populations including four (LD, FR, JM and BE) from southern locations (60 – 62°N) (Figure 6C). The second genetic cluster comprised five cultivars (AK, CO, MB1, UN and UT) and all remaining *C. canadensis* field-collected individuals from more northern latitudes (68 – 69°N) (Figure 6C). These results suggest that cultivated seeds generally represent genetic variation found among wild populations of *C. canadensis* in the Northwest Territories, but seed sources from northern and southern regions are genetically distinct.

Bucharova et al. (2019) and Thomas et al. (2014) aregue that provenances for seed mixtures should be guided by species-specific life history traits, genetic differentiation levels, local adaptation, and ecological goals, which may vary by project (Bucharova et al., 2019; Thomas et al., 2014). The genetic clusters we observed among cultivated and wild populations of *C*. *canadensis* from the Northwest Territories, could enhance provenancing success, whether the goal is to maximize locally adapted gene complexes within regions or increase standing genetic variation by mixing seeds across both regions.

## 5. Conclusion

Genetic studies focusing on ecologically important native taxa like *Calamagrostis* inform conservation and restoration goals. *Calamagrostis* species are valuable for restoration efforts because they can establish and thrive in eroded, acidic and disturbed soils and they play key roles in boreal and arctic ecosystems, such as food sources for elk and bison—two species with high cultural and spiritual value for Canada’s First Nations. Our study highlights the complex genetic ancestry and diversity of three widespread *Calamagrostis* species in the Northwest Territories. Our genome-wide genetic markers provide insights into historical gene flow among species, contemporary hybridization, regional differentiation and genetic differences between wild and commercially cultivated seed sources. We developed a species-specific GT-seq SNP panel with modified analysis pipeline for polyploids to help address a gap in arctic research and inform provenancing to restore or maintain ecosystem resilience in vulnerable northern environments. Ultimately, we hope that our study can contribute towards efforts of fostering ecosystem restoration accountability and effective reconciliation with First Nations.

## Acknowledgements

Seed collections were provided by the Aurora Research Institute (ARI) in the Northwest Territories. Computing resources were provided by BCNet Compute Canada (part of Digital Alliance Canada) and the Center for Advanced Computing at Queen’s University. The authors acknowledge Olivia Bain and Heidi Albers for assistance with data collection and lab work, greenhouse manager Saeid Mobini support with phytotron work. Funding was provided by an NSERC Discovery Grant RGPIN-2023-05811 to RIC and a Northwest Territories Environmental Research Fund to PSH and RIC.

## Data Accessibility Statement

FASTA, VCF and CSV files for ddRAD-seq and GT-seq, downstream analysis scripts and metadata (including latitude and longitude of collections and collection region) and are maintained on GitHub (https://github.com/gomezquijano/Calamagrostis_PPLANTs) with reproducible code available through the Dryad Repository (LINK TBD) based on sequencing data available from NCBI: SUB15805174.

## Benefits-Sharing Statement

This research was conducted in compliance with the Convention on Biological Diversity and the Nagoya Protocol. Seed collections were conducted between 2005-2008 under the licence number 13896 on disturbed areas in the Inuvialuit Settlement Region and east of the Dempster between Inuvik and Tsiigehtchic by members of the Aurora Research Institute (ARI) in the Northwest Territories. The authors are committed to sharing all relevant data and outputs generated from this study with local stakeholders to support future conservation work.

## Author Contributions

MJGQ, PSH, and RIC designed the study based on seed collections and associated metadata provided by PSH and funding managed by PSH and RIC. MJGQ led the data analysis and manuscript writing based on laboratory work and preliminary data analyses by MJGQ, YW, AVL, and CS. ZS and NC provided advanced technical support with key aspects of the library preparation and bioinformatics pipeline.

---

**Supplementary Figure 1.**
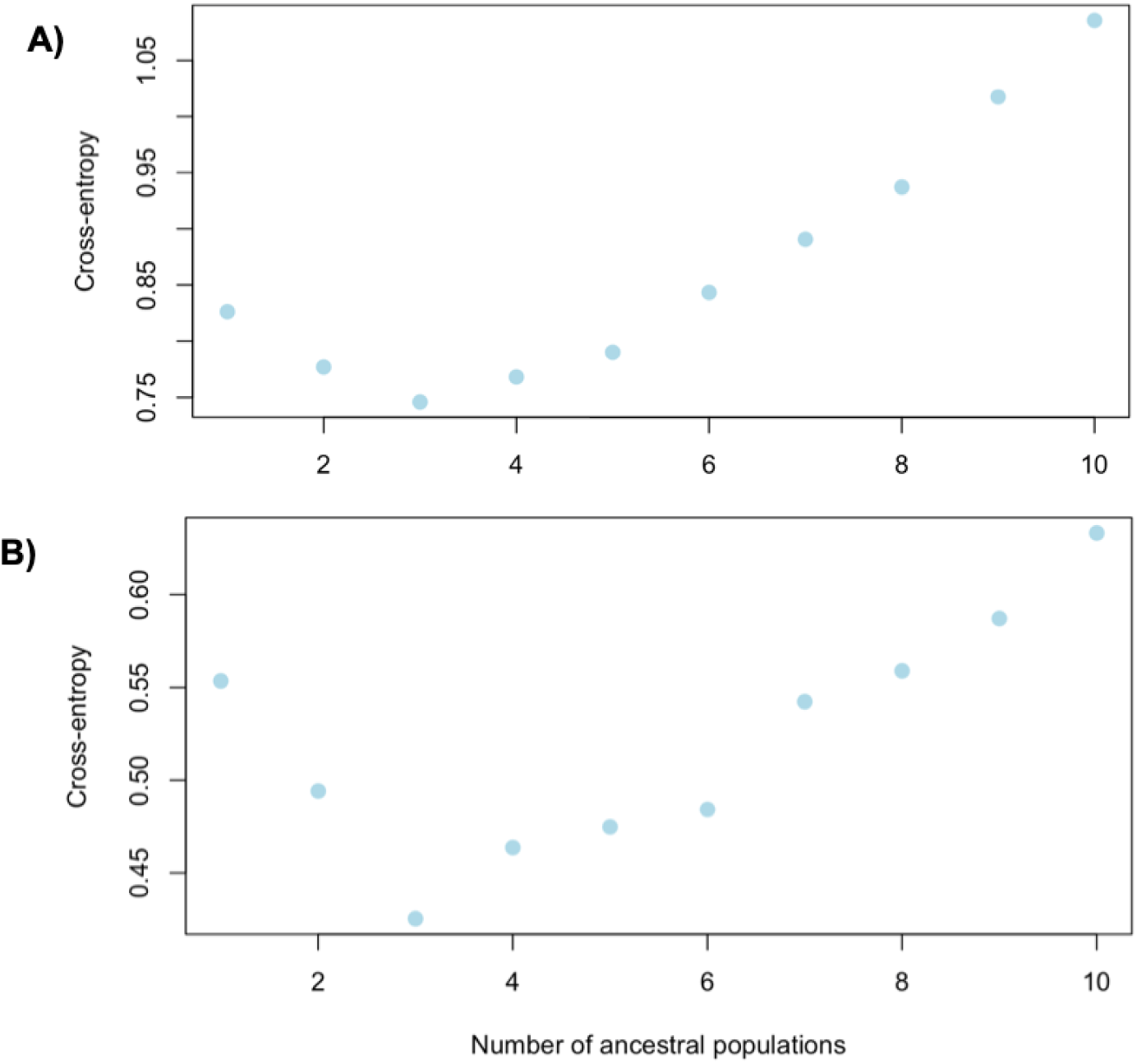
Average cross entropy of five repetitions with 1000 iterations and a regularization parameter of 50 across ten ancestral populations (levels of *K*). Cross entropy values assess predictive performance of the LEA ancestry model for three *Calamagrostis* species using A) ddRAD dataset and B) GT-seq validation dataset. Lowest cross entropy value represents the optimal *K* value.

