## Supplementary figures and images for "Genome-wide SNP diversity in natural and cultivated populations informs restoration ecology with three *Calamagrostis* species from the Northwest Territories of Canada"

### Supplementary Figure 1

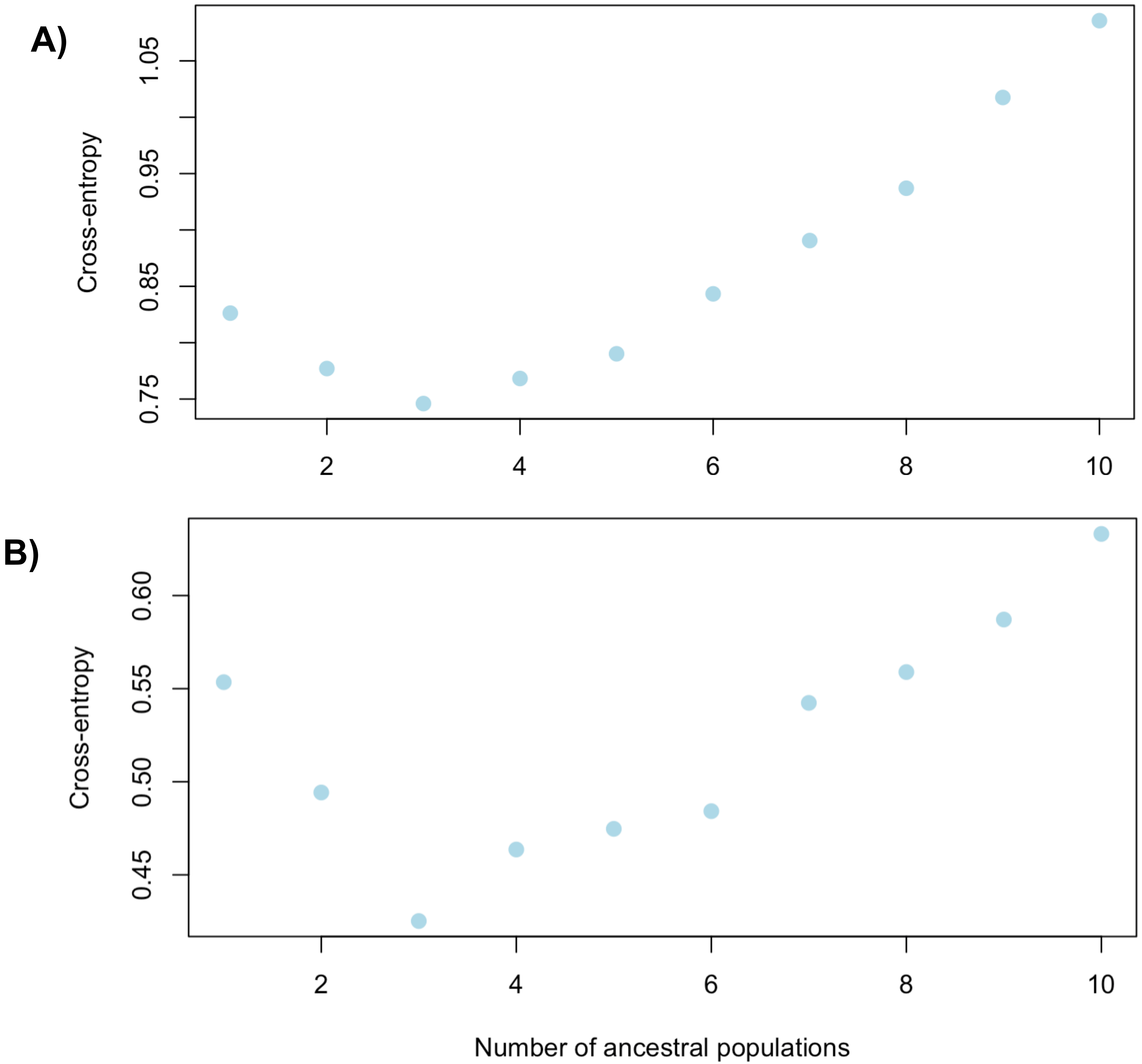
